# Flagellar Motility is Mutagenic

**DOI:** 10.1101/2024.06.21.600093

**Authors:** Souvik Bhattacharyya, Shelby Lopez, Abhyudai Singh, Rasika M. Harshey

## Abstract

Flagella are highly complex rotary molecular machines that enable bacteria to not only migrate to optimal environments but to also promote range expansion, competitiveness, virulence, and antibiotic survival. Flagellar motility is an energy-demanding process, where the sum of its production (biosynthesis) and operation (rotation) costs has been estimated to total ∼10% of the entire energy budget of an *E. coli* cell. The acquisition of such a costly adaptation process is expected to secure short-term benefits by increasing competitiveness and survival, as well as long-term evolutionary fitness gains. While the role of flagellar motility in bacterial survival has been widely reported, its direct influence on the rate of evolution remains unclear. We show here that both production and operation costs contribute to elevated mutation frequencies. Our findings suggest that flagellar movement may be an important player in tuning the rate of bacterial evolution.

---

Assembly and function of bacterial flagella requires more than 50 proteins (1). In *E. coli*, the long external flagellar filament composed of thousands of FliC proteins is attached via a flexible hook to a rod that is stabilized by stationary P and L rings in the outer membrane and connected to a basal inner membrane structure composed of MS and C rings, also referred to as the rotor (Fig. 1A). The top of the C ring is surrounded by multiple stator units composed of MotAB proteins. Proton transit through these units results in rotation of pentameric MotA, which pushes on the C ring, driving rotation of all the connected parts (2). This behemoth of a contraption is ∼20 MDa, excluding the long filament; the filament itself is ∼1.6 GDa (3). The C-ring is crucial for flagellar rotation and for switching rotation direction, enabling ‘run and tumble’ motions that allow chemotaxis (4). Chemotactic signals are generated at a polarly located ‘patch’ composed of thousands of chemoreceptors (MCPs) that detect environmental cues. MCPs control the activity of a linked histidine kinase CheA, which generates chemotactic signals by phosphorylating the response regulator CheY. CheY∼P binding to the C-ring controls rotor switching, this signal terminating upon interaction with CheZ phosphatase (Fig. 1A). A second adaptation circuit composed of CheB and CheR proteins resets the signaling conformation of the MCPs through methylation/demethylation reactions (not shown in Fig. 1A). The cost of production and assembly of all these complexes, which includes ATP-driven export of rod-hook-filament subunits, are estimated to expend ∼5% of the total energy of an *E. coli* cell (5). There is also a high turnover of the flagellin protein FliC because filaments break due to viscous shear (6). The operational cost of rotation, where protons flow from the periplasm through the stators into the cytoplasm, is also reported to be high at ∼5.2% for *E. coli* (5). Each revolution of the rotor requires ∼50 protons per stator unit (5). While the basal structure can spin at the top speed of ∼300 Hz with a single stator unit, assembly of the long external filament increases the load on the basal structure, requiring the engagement of multiple such units (3, 7). Together, flagellar motility can easily take up ∼10% of the total energy budget of the cell (5). Retention of such an energy-guzzling machine must come with long-term benefits because wasteful molecular processes are always under evolutionary scrutiny, and any that impart a fitness disadvantage face elimination over time (8).

**Fig. 1.**
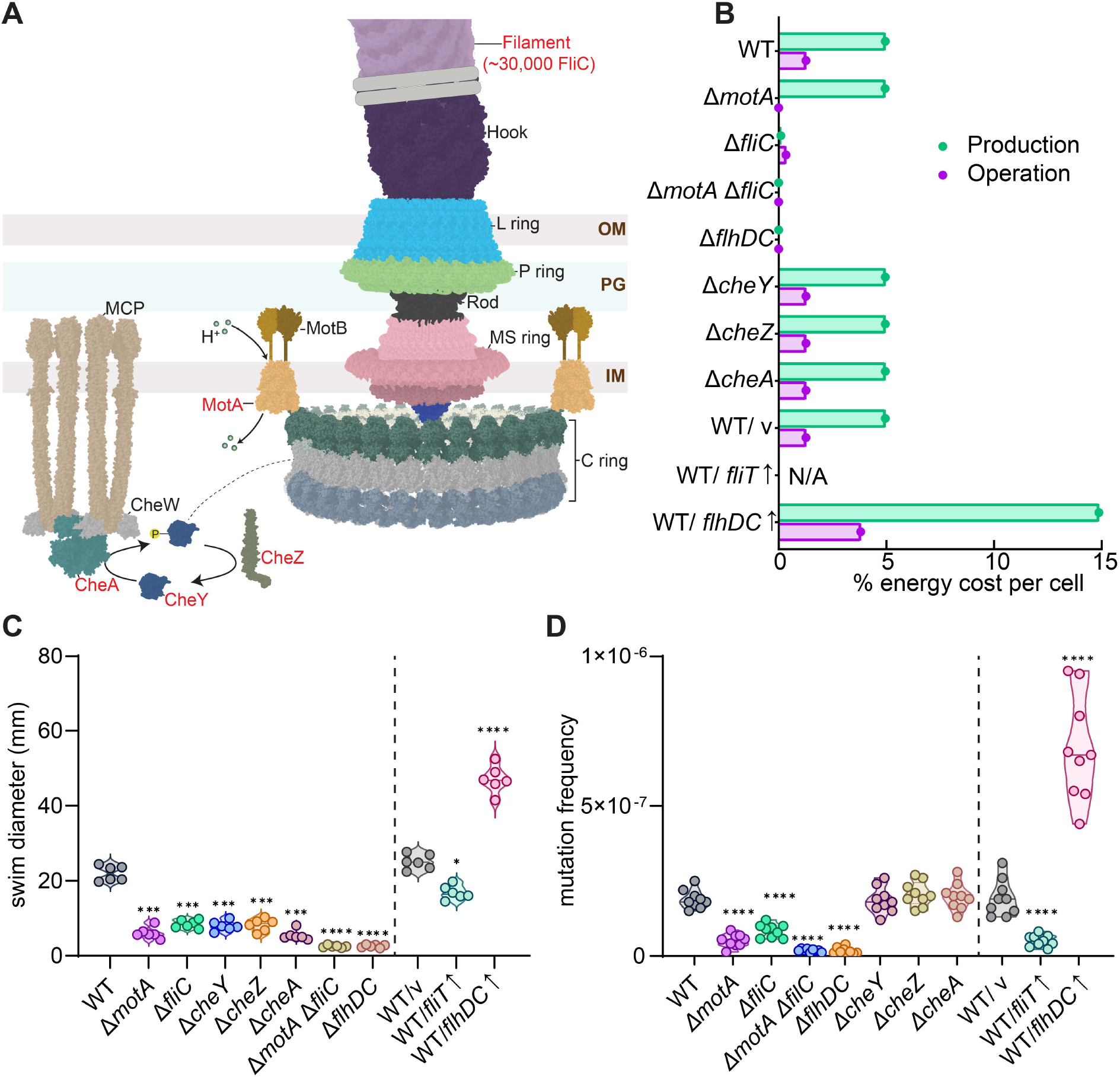
Flagellar motility and mutation frequency are positively correlated. (A) Illustration of the flagellar apparatus and chemotaxis system derived from structures of experimentally resolved components from *E. coli* and related bacteria (drawn to scale). PDB IDs are: 7CGO (*S. enterica* hook, L/ P/MS-rings, rod and export apparatus); 8UOX (*S. enterica* C ring composed of FliG-FliM-FliN); 1UCU (*S. enterica* FliC filament); 6YKM (*C. jejuni* MotAB), 8C5V (*E. coli* chemoreceptor apparatus consisting of MCP, CheY, and CheW); 1F4V (*E. coli* CheY); 1KMI (*E. coli* CheZ). See SI Methods for details. The genes mutated in this study are labeled in red. (B) Operational and production costs of deletion (Δ) and overexpression (*↑*) strains, calculated as per (5) and (7). See SI Methods for calculation details. (C) Swimming motility (6h) of various strains assayed on soft agar (n=6); v, empty vector control for the overexpression strains. (D) Mutation frequencies (MF) of indicated strains measured as Rif^50^-resistant CFUs per unit CFU (n=9). The ±fold changes in MF (in parenthesis) compared to WT are: Δ*motA* (-3.5), Δ*fliC* (-3), Δ*motA*Δ*fliC* (-10), Δ*flhDC* (-10), *fliT↑* (-2), *flhDC↑* (+4.5). All p values were calculated from Mann-Whitney tests by comparing each strain with respective WT controls; ^∗^p < 0.05, ^∗∗^p < 0.01, ^∗∗∗^p < 0.001, ^∗∗∗∗^p < 0.0001.

Efflux pumps are another example of an energy-intensive process, which we recently showed can accelerate evolution in *E. coli* by promoting mutations (9). Such processes generate ROS species that cause oxidative damage to all cellular components including DNA, and hence are mutagenic (10). Cells respond to this damage by deploying ROS scavenging systems (10). However, mutagenic events can be beneficial under specific stress conditions, providing a bet-hedging survival strategy (8, 9). Since flagellar motility is important for survival in diverse stress conditions such as during host colonization (4) and antibiotic survival (9), we posited that it should also promote mutations through the same energy-ROS-mutation nexus. The experiments described below were designed to test this hypothesis.

## Results and Discussion

We first created mutants exhibiting defects in flagellar function (Fig. 1A, red fonts). The flagellar energy expenditure of WT and mutants was recalculated based on a previous report (5) (see SI Methods). Our calculations yielded ∼4.95% and ∼1.27% production and operation costs, respectively, for WT, the latter being different from the previous report (5) because of a recent report that only half of the normal complement of stators are employed during swimming (7); this number is expected to increase when bacteria swim through viscous media or swarm on a surface (11). Deletion of *fliC* would reduce production costs by 97% and deletion of *motA* would reduce operational costs by 100%. Deletion of individual chemotaxis components, however, would have smaller effects (0.002% reduction in Δ*cheA*, for example). Deletion of the flagellar master regulators *flhDC*, or a double deletion of *motA-fliC*, would eliminate the total cost entirely. As expected, deletion of *fliC, motA* or *flhDC* completely abolished motility (Fig. 1C); chemotaxis mutants do not migrate out on motility plates but are otherwise motile. We also tuned expression of the flagellar regulon down or up by either overexpressing (↑) the anti-*flhDC* factor *fliT* (12) or *flhDC* itself (13), decreasing and increasing motility, as expected (Fig. 1C). The mutation frequency (MF) of all these strains was tested by scoring for Rif^50^resistant colonies (14). Loss or diminution of motility decreased MF (Fig. 1D, Δ*motA*, Δ*fliC*, Δ*flhDC, fliT↑*), while increased motility had the opposite effect (Fig. 1D, *flhDC↑*). Disrupting chemotaxis (Δ*cheY/Z/A*) had no effect. Major reduction in both production (Δ*fliC*) or operation (Δ*motA*) costs or both (Δ*motA*Δ*fliC*, Δ*flhDC*) reduced MF significantly (see Fig. 1 legend for fold changes).

To test if the motility-MF correlation is mediated via ROS generated during energy consumption, we used CellROX dye to measure redox levels (9). These levels (Fig. 2A) followed the same trend as that of MF across all strains as seen in Fig. 1D. Fluorescence microscopy confirmed these observations in Δ*motA* and Δ*fliC* strains: both strains had lower ROS levels compared to WT (Fig. 2B). To test these results by a second method, we employed the redox dye 5-cyano-2,3-ditolyl tetrazolium chloride (CTC), which is reduced by electron transport chain activity to form fluorescent CTC-formazan (9). Δ*motA*, Δ*fliC*, Δ*motA*Δ*fliC*, Δ*flhDC*, and *fliT↑* strains all showed a significant decrease in the CTC signal, while *flhDC↑* showed an increase (Fig. 2C). Addition of the antioxidant glutathione (GSH) (15), drastically reduced MF in the *flhDC*↑ strain (Fig. 2D, left), ascertaining the contribution of ROS to the observed MF. (Data in Fig. 2D right, are controls for those in Fig. 1D: complementation of Δ*motA* and Δ*fliC* with plasmids expressing *motA* and *fliC*, restored their MF to WT levels). When the MF values of all tested strains (except WT/ *fliT↑*, for which, quantitative experimental data is yet to be reported) were plotted against calculated energy costs, a linear dependence was obtained (Fig. 2E); the strong correlation coefficient (R^2^=0.9341) indicates that the two variables – MF and the total cost of motility – are highly linked. The nature of the lower and upper bounds of this linearity is a topic for future investigations.

**Fig. 2.**
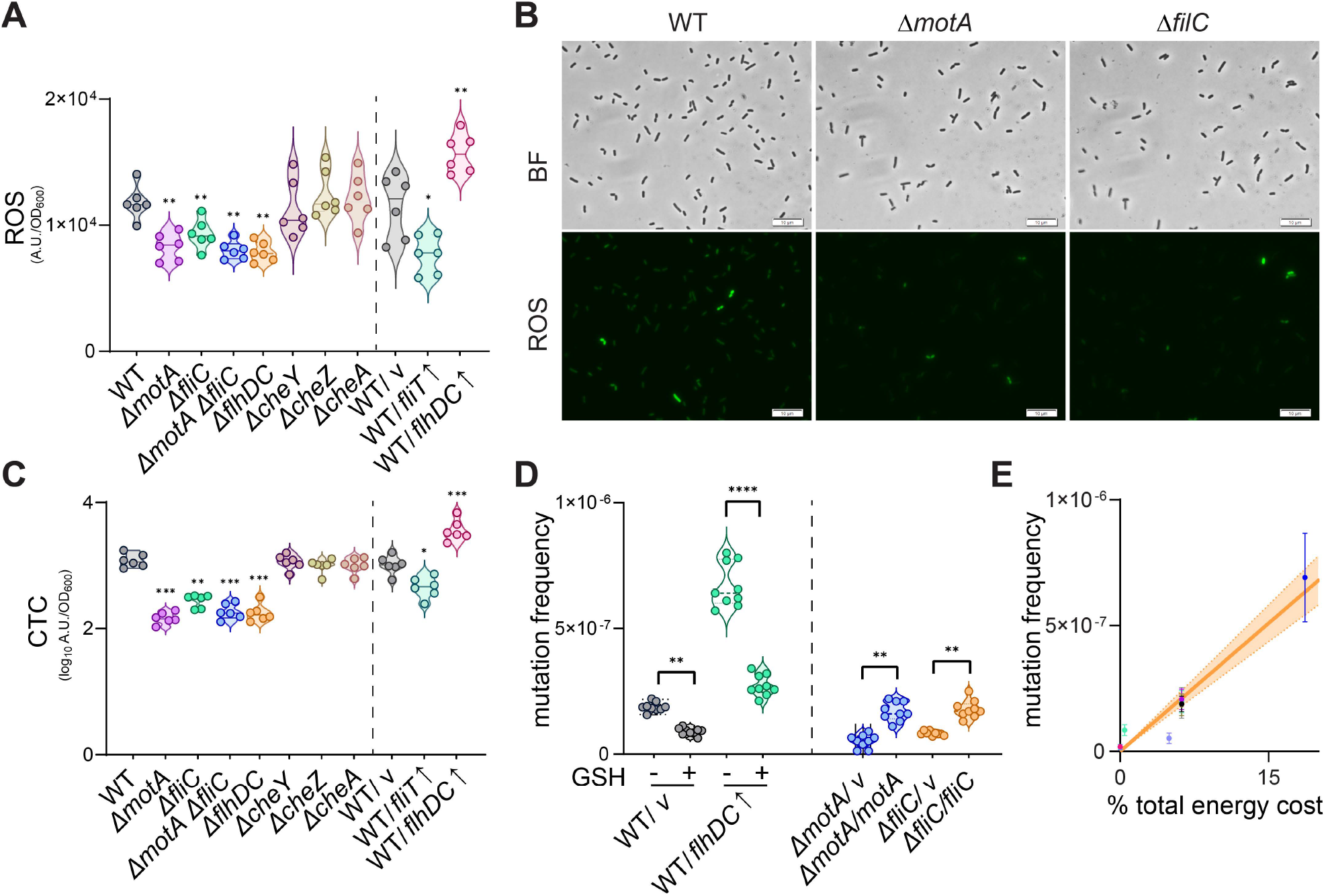
Mutagenicity of flagellar motility is ROS derived. (A) ROS levels in different strains using CellROX redox sensor (n=6 for each strain). p values were calculated from Mann-Whitney tests. (B) Brightfield (BF) and ROS green fluorescence images of different strains. Mid-log cells were used for staining with Cell-ROX dye and imaged. (C) Energy metabolism was estimated by staining the cells with CTC dye (n=6 for each strain). The p values were calculated from Wilcoxon rank-sum tests. (D) MF was measured as in Fig. 1D; ±GSH, 50 mM glutathione. The right panel shows restoration of MF in Δ*motA* and Δ*fliC* strains after complementation with *motA* and *fliC* expressed from plasmids. p values were calculated from Wilcoxon rank sum tests (left) or Mann-Whitney tests (right); ^∗^p < 0.05, ^∗∗^p < 0.01, ^∗∗∗^p < 0.001, ^∗∗∗∗^p < 0.0001. (E) Correlation of MF and flagellar energy cost of all strains (except WT/ *fliT↑*) using data shown in Fig. 1D and 1B respectively; R^2^=0.9341.

In summary, our data provide strong experimental support for the hypothesis that flagellar motility incurs mutational costs through the energy-ROS-mutation nexus. Bacteria likely exploit this nexus to transition between high and low MF states, depending on their environmental niche. This is the second example of a high energy-demand / high-mutation connection (9), and the reason why both flagellar motility and efflux pumps are tightly controlled by a vast network of regulators. Our findings beg the question: did the innovation of a functional flagellum accelerate the rate of bacterial evolution?

## Acknowledgements

We thank lab members Nabin Bhattarai and Yuki Hyodo for generating Δ*cheA* and Δ*fliC-motA* mutants respectively. This work was supported by NIH grants GM118085 (NIGMS) and AI158295 (NIAID) to R.M.H., and R35GM148351 (NIGMS) to A.S. S.B. is a Provost’s Early Career Fellow.

## Author Contributions

S.B. conceived the project; S.B. & R.M.H designed the experiments; S.B. & S.L. performed the experiments; S.B., S.L., and A.S. performed the energy calculations; S.B., S.L, & R.M.H analyzed and interpreted experimental data; S.B. performed statistics; A.S. performed modeling; S.B. & R.M.H wrote the paper.

## Supplementary Information

The SI file contains SI Methods and SI References.

