## Supplementary material for "Flagellar Motility is Mutagenic": SI

### SI Methods

**Strains, plasmids, media, and genetics**

Strains were propagated in LB broth (10 g/L tryptone, 5 g/L yeast extract, 5 g/L NaCl) or on 1.6% Bacto agar plates with the following antibiotics for marker selection as applicable – Kan (Kanamycin) 25 µg/ml, Amp (Ampicillin) 100 µg/ml, Cam (Chloramphenicol) 30 µg/ml. *motA*, *fliC*, *cheY*, and *cheZ* deletion strains were previously created in the lab (1-3) using the one step deletion method, followed by removal of the Kan cassette via pCP20 method (4). ∆*cheA* and ∆*motA*∆*fliC* strains were generated similarly in this study by deleting *cheA* and *motA* genes in WT and ∆*motA* strains, respectively. The ∆*flhDC* strain was previously described (1). All deletions were confirmed by colony PCR. Overexpression of *flhDC*, *motA*, and *fliC* was achieved with 0.5% arabinose induction from the pBAD33 plasmid. Overexpression of *fliT* was achieved with 1 mM IPTG induction from pCA24N plasmid from the ASKA collection (5).

**Energy calculations**

To determine the energy cost of the *E. coli* flagellum we used the calculations of Schavemaker & Lynch (see Table 1 and its source data in (6)), where the production cost for WT was calculated to be $7.88 \times 10^{8}$ ATP per cell, which is approximately 5% of the total cell energy cost of $1.59 \times 10^{10}$ ATP per cell. These production costs were assumed to be the same for *motA*, *cheA*, *cheY*, and *cheZ* deletion strains. Since close to 98% of the production cost is the synthesis of the filament, the production cost for *fliC* deletion strains was taken to be $0.17 \times 10^{8}$ ATP per cell (6).

For computing the operation cost of the *E. coli* flagellum, we again followed Schavemaker & Lynch (6). While the motor can have a maximum of 11 torque-generating stator units, only half this number was reported for *E. coli* during swimming (see Fig. 1 in (7)). Based on these more recent measurements, the operation costs were calculated as in (6), but assuming an average of 5.5 stators and a flagellum rotation rate of 100 Hz (Fig. 5 in (7)). As per this calculation, we obtained the operation cost for WT to be $2.2 \times 10^{8}$ ATP per cell or 1.27% of the total energy cost, and this cost was assumed to be the same for *cheA*, *cheY*, and *cheZ* deletion strains. For the *fliC* deletion strain, we considered only 1 stator unit per motor given the reduced load. The operation cost was zero for *motA* deletion strains, and both operation and production costs were zero for ∆*flhDC* strain. In the case of *flhDC* overexpression, both costs were scaled by three times following the previously reported experimental data of flagellar number in that strain (1).

**Swim assay**

From mid-log phase cultures (0.6 $OD_{600}$), 4 µL of cell suspension were stabbed at the center of 0.3% LB agar plates, followed by incubation at 37°C for 6h.

**Mutation frequency**

For each strain, 9 overnight cultures were sub-cultured (0.01% inoculum) into 5 ml of fresh media and grown till 0.6 OD. Then, 1 ml culture (adjusted to ~$OD_{600}$=1) was plated on 50 µg/ml Rifampicin (Rif) LB agar (1.5% w/v) plates and incubated at 37°C for 24 hours. From the same cultures, CFUs were counted on LB agar plates (1.5%) without antibiotic, using a serial dilution method. The resistant colonies were counted on Rif plates. MFs were calculated as Rif-resistant colonies/CFU. To avoid jackpot mutation events in a population, the results were considered to be usable only when the median values of MFs were within 95% CI from the same-day data (8, 9). Since Rif is light-sensitive (10, 11), all Rif plate work was done in near-dark conditions with no direct light inside a laminar flow. Given the complexity of the genetic organization of flagellar operons (12), we took an abundance of precaution to eliminate polar effects in our MF data by complementing the deleted genes in the Δ*motA* and Δ*fliC* strains.

**ROS assay**

Cells were stained using CellROX Green Reagent Kit (Thermo) following manufacturer's protocols. The CellROX dye, which is weakly fluorescent in its reduced state, increases fluorescence upon oxidation (Ex, 485 nm; Em, 520 nm). Mid-log phase cells (0.6 $OD_{600}$) were incubated with the dye (500:1) for 15 min in the dark at room temperature. Fluorescence was 1) measured by a microplate assay using Spectramax m3 reader as arbitrary units or A.U.= $A_{520}$ / $OD_{600}$, and 2) visualized under a light microscope (BX53F; Olympus, Tokyo, Japan) with either brightfield or GFP fluorescent channel (Ex, 460–480 nm; Em 495–540 nm, 100 ms exposure time). Images were acquired via cellSens software (v1.6) using a 100X objective and XM10 CCD camera (Olympus, Tokyo, Japan).

**CTC assay**

To measure active respiration using 5-Cyano-2,3-ditolyl tetrazolium chloride or CTC (13), 100 µl of mid-log phase cells (0.6 $OD_{600}$) were treated with 2mM CTC (Sigma) and shaken at 200 rpm for 10 min at 37 °C. Fluorescence was measured (Ex, 485 nm; Em, 630 nm) by a microplate assay using Spectramax m3 reader as $Log_{10}$A.U.= ($A_{630}$ / $OD_{600}$).

### Statistics and visualization

Statistical analyses were performed using Prizm v10 (Graphpad) and Microsoft Excel. The data were visualized in Prism.

### Flagellar model

To create the diagram shown in Fig 1A, the indicated pdb files were downloaded from RCSB protein data bank, visualized in ChimeraX (14), followed by importing the png files to Adobe Illustrator. The diagram was given a pastel look by layering vector outlines of each component on top of the original image. The flagellar structures and the membrane components were drawn to scale using previous literature ((15) and (16, 17) respectively).
